# Models as games: a novel approach for ‘gamesourcing’ parameter data and communicating complex models

**DOI:** 10.1101/2021.09.23.461497

**Authors:** Jeroen Minderman, A. Bradley Duthie, Isabel L. Jones, Laura Thomas-Walters, Adrian Bach, Matthew Nuttall, Nils Bunnefeld

## Abstract

1. Models have become indispensable tools in conservation science in the face of increasingly rapid loss of biodiversity through anthropogenic habitat loss and natural resource exploitation. In addition to their ecological components, accurately representing human decision-making processes in such models is vital to maximise their utility. This can be problematic as modelling complexity increases, making them challenging to communicate and parameterise.
2. Games have a long history of being used as science communication tools, but are less widely used as data collection tools, particularly in videogame form. We propose a novel approach to (1) aid communication of complex social-ecological models, and (2) “gamesource” human decision-making data, by explicitly casting an existing modelling framework as an interactive videogame.
3. We present players with a natural resource management game as a front-end to a social-ecological modelling framework (Generalised Management Strategy Evaluation, GMSE). Players’ actions replace a model algorithm making management decisions about a population of wild animals, which graze on crops and can thus lower agricultural yield. A number of non-player agents (farmers) respond through modelled algorithms to the player’s management, taking actions that may affect their crop yield as well as the animal population. Players are asked to set their own management goal (e.g. maintain the animal population at a certain level or improve yield) and make decisions accordingly. Trial players were also asked to provide any feedback on both gameplay and purpose.
4. We demonstrate the utility of this approach by collecting and analysing game play data from a sample of trial plays, in which we systematically vary two model parameters, and allowing trial players to interact with the model through the game interface. As an illustration, we show how variations in land ownership and the number of farmers in the system affects decision-making patterns as well as population trajectories (extinction probabilities).
5. We discuss the potential and limitations of this model-game approach in the light of trial player feedback received. In particular, we highlight how a common concern about the game framework (perceived lack of “realism” or relevance to a specific context) are actually criticisms of the underlying model, as opposed to the game itself. This further highlights both the parallels between games and models, as well as the utility of model-games to aid in communicating complex models. We conclude that videogames may be an effective tool for conservation and natural resource management, and that although they provide a promising means to collect data on human decision-making, it is vital to carefully consider both external validity and potential biases when doing so.

## 2 Introduction

In recent years, the use and application of models1 has become widespread and indispensable in conservation science, ranging from demonstrating the likely effects of climate change on biodiversity (IPCC 2021) to supporting the understanding of fundamental processes in natural resource management (e.g. Schlüter et al. 2012; Fryxell et al. 2010; Cusack et al. 2020). Given the continued rapid global loss of biodiversity (Ceballos et al. 2015; Ceballos, Ehrlich, and Dirzo 2017), understanding the mechanisms and consequences of such loss is vital for long-term sustainability. Although a number of drivers of biodiversity loss have been identified (e.g. Maxwell et al. 2016), one of the most prevalent and widespread is human exploitation of habitats and natural resources, both directly (e.g. through hunting or habitat loss to agriculture) or indirectly (e.g. through international trade in natural resources) (Wilting et al. 2017). Because resource use is fundamentally driven by economic and social processes, accurately predicting future changes is reliant as much on understanding human behaviour and decision-making (Milner-Gulland 2012; Schlüter et al. 2012) as it is on understanding resource dynamics themselves. Thus, the development of social-ecological models in which natural resource dynamics and human decision making interact is becoming increasingly urgent.

Cutting-edge modelling approaches have made significant progress towards this goal. For example, Orach, Duit, and Schlüter (2020) used an agent-based model to show how coalitions of interest groups can stabilise natural resource dynamics, whereas Cusack et al. (2020) used a novel agent-based modelling framework (Duthie et al. 2018) to assess the effect of lobbying on species extinction risk. Although such efforts represent significant progress in modelling complex social-ecological systems, their increased complexity poses two interlinked challenges. First, models are often difficult to communicate clearly to non-specialist audiences, and this challenge increases with model complexity (Grimm et al. 2006). This is particularly important for models of resource use in social-ecological systems, as they are often specifically intended for use by managers or stakeholders who may not have the required technical expertise. Much has been said about improving the uptake of models in such settings (e.g. Bunnefeld, Nicholson, and Milner-Gulland 2015; Addison et al. 2013; Schuwirth et al. 2019; Will et al. 2021), and detailed documentation of the purpose, organisation and predictions has been highlighted as particularly important (Grimm et al. 2020). Even so, frequently the evidence for practical uptake of many models is limited (Addison et al. 2013; Bunnefeld, Nicholson, and Milner-Gulland 2015; Zasada et al. 2017). Second, their complexity implies the need for extensive data to parameterise them effectively. In terms of social-ecological systems, while data to parameterise the ecological component are often relatively easily available, the human decision-making components are often based on limited theory and lack a general empirical basis (Groeneveld et al. 2017; Schwarz et al. 2020). Not only may this lead to limited predictive power, but stakeholders may also be unwilling to accept model results that they perceive as lacking an empirical basis (cf. model “quality” as in Kolkman et al. 2016). To maximise the adoption of complex social-ecological models as management tools, both appropriate representation of human decision-making, and effective communication, are therefore key.

Games have a long history of use in research (Sandbrook, Adams, and Monteferri 2015; Chabris 2017; Redpath et al. 2018), including as tools to aid the communication of complex ideas and processes to non-specialists (Garcia, Dray, and Waeber 2016; Tan et al. 2018), with recent work starting to leverage online and video games (Oultram 2013; Pérez and Guzmán-Duque 2014; Fjaellingsdal and Kloeckner 2019; Crowley, Silk, and Crowley). Given this long history, it is striking that the parallels between games, in particular videogames, and models are not recognised more widely. All models are abstract representations of environments, actors and relationships, with inputs (parameters) and outputs (predictions or inferences). Similarly, all games present a player with an environment in a given state (parameters), including one or more actors, who can take actions (inputs) to affect the environment for a given effect (outputs). It is worth stressing that every game has an underlying model that defines the state of the environment, relationships between objects in this environment, and inputs and outputs available to the player. However, while games are by definition designed with player interaction in mind, models rarely have user-facing or even user-friendly interfaces, and running or adapting them to specific circumstances usually relies on technical expertise. Casting models as games provides an opportunity to effectively improve the communication and understandability of even relatively complex models. Inputs and outputs may be presented in a visual way and adapted depending on the type of audience, and both potential applications and limitations of the model can be demonstrated effectively.

In addition, presenting a model as a game provides an opportunity to empirically collect data on how stakeholders make decisions in the modelled environment. Games have already been widely used for data collection to answer specific questions (e.g. Meinzen-Dick et al. 2016; Villamor and Badmos 2016; Rakotonarivo, Bell, et al. 2021; Rakotonarivo, Jones, et al. 2021) about what affects decision-making in social-ecological systems. Another potential application of presenting model as games, which warrants further exploration, is using in-game decisions as a “big data” source to improve the parameterisation of the underlying model itself. Many existing models represent human decision-making by relatively crude algorithms (e.g. fully rational utility maximisation) despite widespread recognition that this does not reflect real-world decision-making (Groeneveld et al. 2017). By presenting real-world stakeholders with in-game decisions that would otherwise be taken by a predefined algorithm, large data sets of actions and outcomes may be collected. Given a large enough sample of players and in-game conditions, such data might then be used to train decision-making algorithms that better reflect human decision-making in natural resource management2. Although this “gamesourcing” or “Gamorithm” (Sipper and Moore 2020) approach has already been widely used in a number of other fields (from crowdsourcing accurate protein-structure models to classifying fluorescence microscopy images, Khatib et al. 2011; Sullivan et al. 2018), it remains rare in conservation science (but see van den Bergh et al. 2021). Thus, model-games can be considered “virtual laboratories” (Duthie et al. 2021) to not only test specific hypotheses or predictions, but potentially also as an effective method to source data to parameterise the underlying models based on in-game decisions by real humans.

We aim to illustrate the potential for this model-game approach, both in terms of aiding model communication as well data collection for improved parameterisation, by introducing Animal&Farm (A&F). We developed A&F as a simple interactive game front-end for a complex social-ecological modelling framework (GMSE), in which the player acts as the manager of a virtual environment in which a population of wild grazing animals (the natural resource) may adversely affect farming yield, with farmers acting to maximise their yield and potentially hunting or deterring (through scaring) the animals. We argue that that by acting as an interface between users (i.e. players) and complex underlying models with many components and assumptions, games can simultaneously (1) aid the communication and useability of the underlying model and (2) can be used to gather data to improve the parameterisation of such models. We first briefly summarise the underlying modelling framework, its potential and limitations. Second, we describe both the structure of A&F itself as well as its database back-end. Third, we outline how this approach may be used to collect data on player decision-making in simulated *in silico* experiments, and present some example results of doing so; noting that these findings are intended as illustrative only. Finally, using test player feedback as a foundation, we discuss both the limitations of this approach as well as its wider potential.

## 3 Outline of approach

A&F is available to play online

A&F consists of two main components; (1) the underlying model(s) describing the wild grazing animal (“resource”) population dynamics, estimates of this population through an observation process, and farmer actions, which are all implemented using the GMSE framework as described below; and (2) the game interface for the underlying model, which allows the player to set management actions (specifically, costs for farmer actions) that would otherwise be determined by the management model in the default GMSE set up.

### 3.1 Underlying model: GMSE

We used GMSE to model the social-ecological system underlying A&F. The GMSE R package (https://cran.r-project.org/web/packages/GMSE/index.html) was designed as a flexible solution for parameterising systems that model the management, observation, exploitation and population dynamics of a natural resource (e.g. a population of hunted wildlife). In this section, we summarise the basic functionality of GMSE as relevant to the present manuscript; for a full description see Duthie et al. (2018) and Nilsson et al. (2021) (the latter containing an appendix with the full ODD model description).

#### 3.1.1 Basic introduction of GMSE principles and structures

GMSE is an agent-based modelling framework consisting of four sequential submodels (Figure 1a) with three types of agents:

1. The **resource model**, consisting of □ individual animal-agents (hereafter referred to as “animals”) moving on a landscape of □ x □ cells.
2. The **observation model** which represents the process of observations (including a degree of uncertainty) of the animal population.
3. The **manager model** consists of a *single* agent (hereafter referred to as the “manager”) which uses the observation of the animal population to make management decisions, affecting the permissiveness of actions for agents in the user model (below).
4. The **user model**, consisting of □ individual agents, which in the current context represent land-owning farmers. Thus, we refer to these agents as “farmers” in the remainder of this manuscript, but note that we use the term “user model” to refer to the general submodel containing these agents to maintain consistency with the term used in the GMSE documentation (Duthie et al. 2018). Each farmer owns a given proportion of the landscape (this may or may not be an equal distribution, see below).

**Figure 1.**
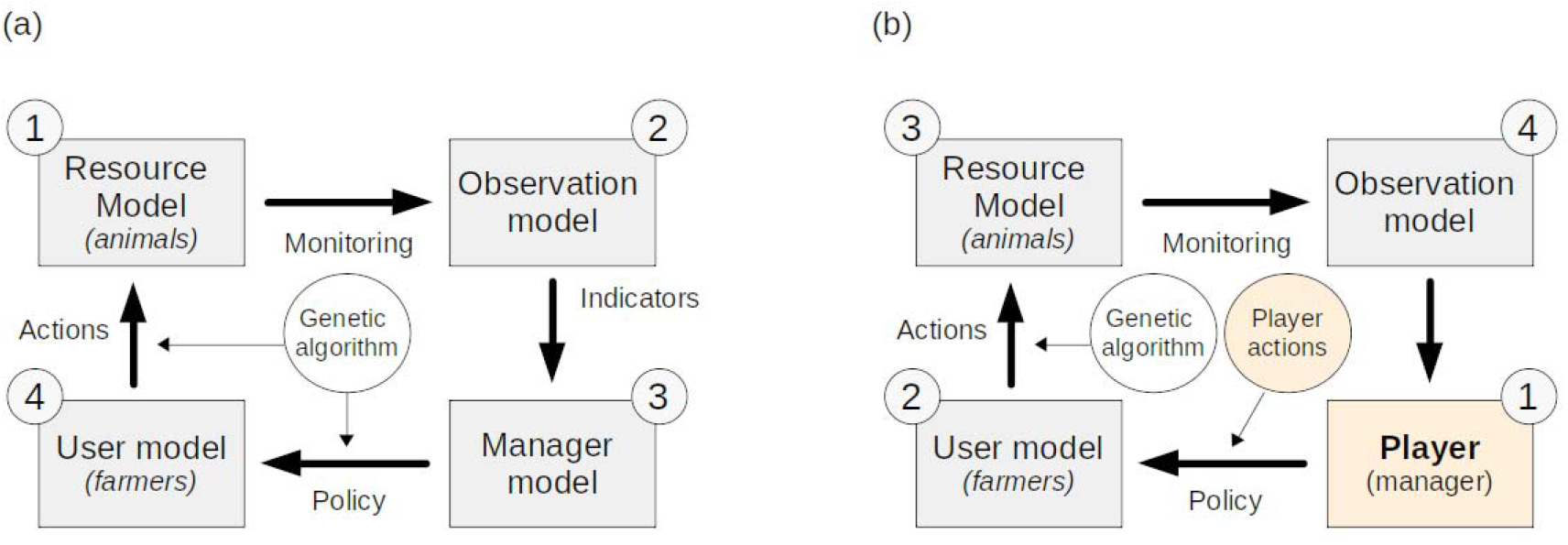
The basic structure of (a) the GMSE modelling framework and its default order of operations with the genetic algorithm (GA) modelling the decision-making process of both users and manager, and (b) the adaptation of the GMSE framework to accommodate the model-game approach presented here. The grey boxes represent the various GMSE submodels, with the arrows representing the process links between them. The yellow boxes and circles are the adapted components in the model-game adaptation, with player interaction replacing the manager model in GMSE, and the underlying GA for the manager - the GA is still used to make user decisions. Grey circles indicate the order of operations in each.

In each GMSE time step, both the manager and all farmers are allocated a (fixed) budget. In GMSE terms, “budget” should be interpreted as a budget of general “resource”; conceptually this may be interpreted as a financial budget, time, materials, or a combination thereof. Farmers may allocate their budget to taking one of several potential actions on their land (here: hunting animals, scaring animals off their land, or tending crops). Both former actions are common in the management and control of grazing animals on croplands (e.g. grazing migratory wildfowl, (Nilsson et al. 2016, 2021), with scaring for example including the use of acoustic deterrents.

The goal for the manager is to maintain the animal population at a desired level (the management target, normally set externally as a model parameter). The manager does so by controlling the cost for farmer actions in the following time step. For example, a higher cost for hunting will decrease the number of animals hunted by farmers leading to population growth, and a lower cost for scaring will increase the number of farmers choosing scaring as an action.

Farmers aim to maximise agricultural yield from their land. By default, yield equals 1 per landscape cell owned per time step, but this may be decreased by the presence of grazing animals in a cell, and/or increased through tending crops. The rates of increase or decrease in yield through grazing and tending crops respectively are controllable in GMSE but kept as constant rates in the current A&F implementation, see Duthie et al. (2018) and parameter references for `res_consume` and `tend_crop_yld` in the `gmse()` function here: https://cran.r-project.org/web/packages/GMSE/GMSE.pdf. Thus, although their objective does not directly relate to the animals, farmers have an incentive to control the number of animals on their land to minimise potential negative effects on their yield. They can do this by allocating budget to hunting or scaring animals. The former reduces the number of animals present in the landscape, while the latter has a certain probability of moving an animal away from the farmers’ land, for the duration of the time step. The relative expected efficacy of the three possible actions (hunting, scaring or tending crops) depends on the number of animals on their land, and the cost of hunting and scaring set by the manager. Farmers can only take actions on land that they own.

By default, costs for farmer actions as set by the manager and actions taken by the farmers are chosen using a genetic algorithm (GA), a heuristic optimisation algorithm that models the choice of decision by mimicking the process of evolution by natural selection: a large number of possible decisions are iteratively compared by assessing their outcome, with the decision that maximises a given utility function (yield for farmers, and minimising distance to population target for the manager, see Duthie et al. 2018) identified as the “fittest” (Hamblin 2013). The GA is run separately for each agent (manager and each user) in each time step.

In the default resource (animal) model in GMSE, the animal population is modelled with a form of logistic growth, with a small amount of random mortality added per time step and death caused by hunting; for more detail see below and Duthie et al. (2018). In each time step, each animal moves a given distance in a random direction, and feeds from the cell in which it is present. In the current model, neither movement nor population growth rate is affected by agricultural yield.

It is worthwhile stressing that in the current GMSE implementation, using the GA, both agent types (farmers and the manager) have only a single goal they each aim for. Farmers aim to maximise their yield, whereas the manager aims to minimise deviation from a given population target - neither can balance multiple competing objectives. This is unlikely to be reflective of real conservation scenarios, where it is common for conservation managers to at least recognise other aims, if not take these explicitly into account when setting policy, and other stakeholders in the system (e.g. farmers) commonly having some interest in conservation objectives (Redpath et al. 2017; Bunnefeld, Nicholson, and Milner-Gulland 2015). Human decision-making in such scenarios is inevitably about balancing these different objectives, but parameterising algorithms that mimic such processes without reference to empirical data is very challenging (Constantino et al. 2021; Dobson et al. 2019). Addressing this issue was a key motivation for the development of the model-game approach presented here.

### 3.2 Animal & Farm

#### 3.2.1 Structure as relating to GMSE

In the default implementation of GMSE v0.7.0.0, a single time □ step consists of a call to the resource model, observation model, management model and user model, in that specific order; in other words, a time step ends after farmer actions have been chosen (by the GA) and implemented (Figure 1a). To allow players to assess the environment and interactively choose management actions, A&F uses a modified version of GMSE implemented in R version 4.1.1 (2021-08-10), the code for which is freely available: https://github.com/ConFooBio/gmse/tree/man_control. In this version, the management model is replaced by player inputs, and the order of operations is altered to accommodate this. Further details are given in S1.

The current GMSE parameter values used by A&F largely reflect default parameter values in GMSE. This is a purely pragmatic choice: because we are not modelling a specific system here, and instead aim only to illustrate the use of the A&F platform in general terms, the specific parameter values given below and in Table S2 should be interpreted as examples: we emphasise that all these parameters are expected to be modified as appropriate for specific GMSE and A&F applications.

The example parameterisation used here simulates a landscape of 100×100 cells, divided into farms owned by 4-12 farmers (the precise number and land distribution is randomly varied per session, see 4.2 below). Farmers can take three possible actions: tending crops, hunting (culling) animals, or scaring animals off their land. All submodels used in A&F are currently the default GMSE models (see S1), with the exception of the management model in time steps □ > *5* where the player assumes control over the management decisions (see below). We only give brief details on GMSE itself here, for full details and descriptions of all models, see Duthie et al. (2018) and Nilsson et al. (2021). The **animal population** model uses the logistic growth form with □_□_ = *1000*, □ = *0.3* and □ = *5000*, meaning that in the absence of any management the population will increase from the initial population size (1000) to carrying capacity (5000). The **observation model** uses the default GMSE model (density-based sampling of a subset of the environment); the manager can only base decisions on the *observed* number of animals (and thus population trajectory plots in the game interface reflect observations only, which are subject to an unknown - to the player - level of uncertainty). Both the **management model** (in the initialisation steps) and **user model** use the genetic algorithm with default parameter settings. Farmer budgets are set to 1500 units per time step, manager budgets to 1000 units (both for the initial 5 time steps and the subsequent game play; see 3.1.1 above for notes on the conceptualisation of “budget”). Farmers aim to maximise yield from their land; their annual budget is reset each year and is unaffected by yield. Yield is positively affected by tending crops and may be negatively affected by the presence of grazing wild animals - thus hunting or scaring may offset any potentially negative effects on yield.

Each subsequent A&F time step consists of (1) player input, taking the place of the default management model, in which the player can assess the environment using outputs provided (see below) and choose management actions (costs for farmer actions), and (2) a modified GMSE time step (following player confirmation of their inputs), including sequential calls to the default user, resource and observation models (`gmse_apply_UROM()`) (Figure 1b).

#### 3.2.2 Player interface

The player interface for A&F is a web application coded in R, using `Shiny` (1.6.0), and packages `shinyjs` (2.0.0), `shinyBS` (0.61), and `waiter` (0.2.2).

On starting a new game session, the player is presented with a series of introductory screens explaining the background, flow, and objective of the game, after which they are asked to enter a player name, which is stored and used to show player scores as the end of a session, compared to previous sessions by other players.

The main game screen consists of four components (Figure 2). First, a trajectory plot (Figure 2a) showing (1) observed animal population numbers and (2) agricultural yield for each farmer in each time step, up to time □ (at the start of the game, this will show five observations from the initialisation steps described above). Agricultural yield is expressed as a % of “maximum unaffected yield,” i.e. yield in the absence of damage from wildlife or investment in tending crops. Second, a plot of the landscape (Figure 2b) showing the distribution of farm ownership as well as the position of animals at time □. Third, a bar plot of the number of actions taken by each farmer at time □ (Figure 2c). Fourth, a report of the current management budget available (not allocated), player scores (see 3.2.3 below), and player inputs (Figure 2d). The player (manager) inputs consist of two sliders, setting the cost for two out of the three actions available3 to farmers in time □ + *1*: killing animals (presented as the cost of a hunting licence) and scaring animals off their land (presented as the cost of a scaring licence). Management budget allocated to one cost cannot be allocated to the other, and any remaining budget is not rolled over to the next time step. The third action available to farmers (tending crops) cannot be directly4 affected by the manager (player), so no input is available for it.

**Figure 2.**
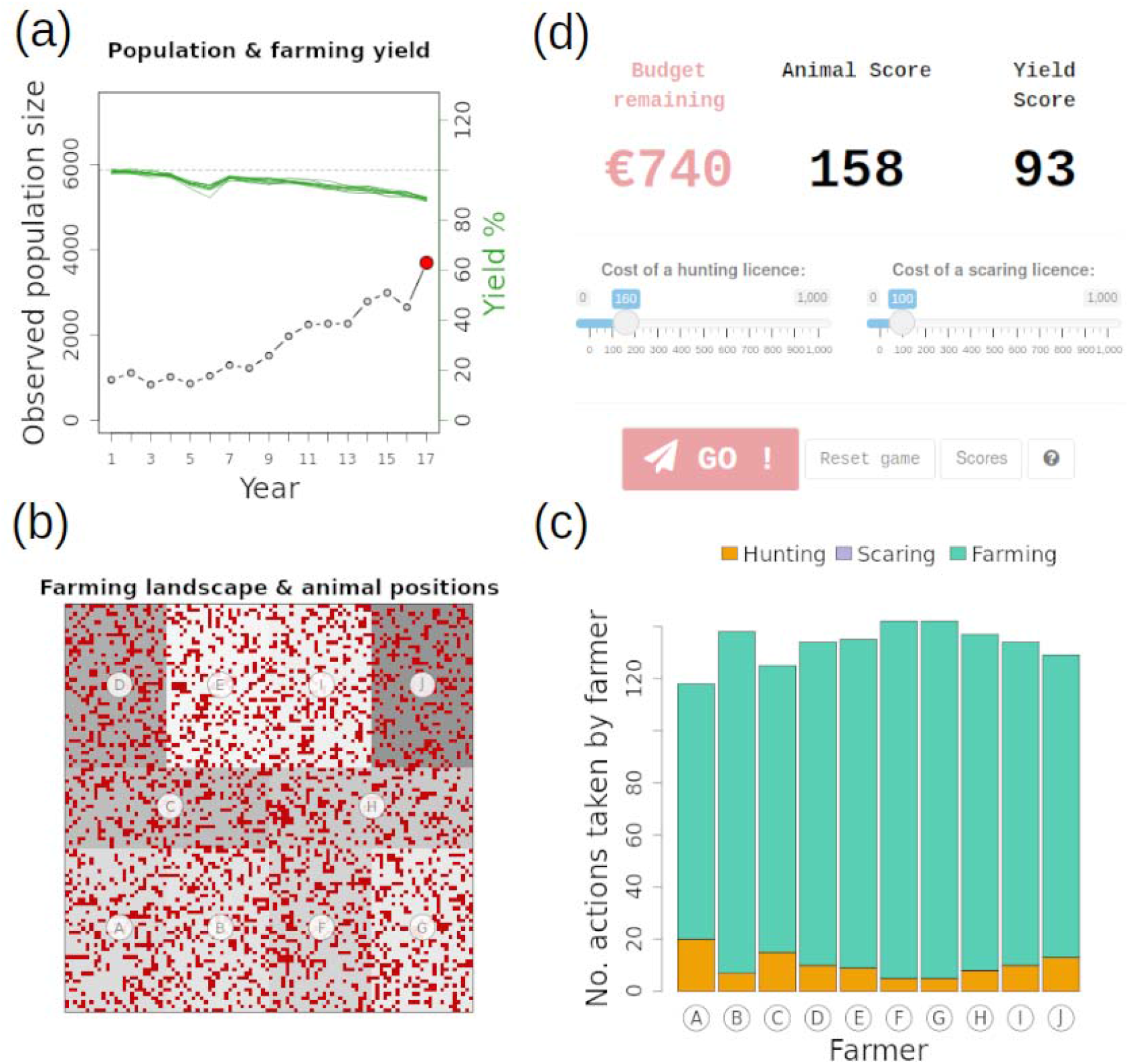
The ‘Animal and Farm’ main game interface, showing (a) the animal (resource) population trajectory and yield per farmer, (b) the farming landscape with animal positions as red dots and farm ownership indicated by the grey shades, (c) actions taken by farmers in the previous game round, and (d) player inputs including a budget report and costs set for actions.

The game progresses to the next time step □ + *1* once the player confirms their choice of cost inputs. At this point (1), the user, resource and observation models are run using the updated action costs set by the player, (2) selected environment state data are stored in the database (See 3.2.3 below), and (3) trajectory, landscape and action plots are updated, and budget allocation is reset. The current implementation of A&F continues for a maximum of 20 time steps (following the initial five) at which point the game session is ended and the player is presented with a scoreboard. If the resource population reaches extinction in any given time step, the game session is also terminated.

#### 3.2.3 Game objective, scores and scoreboard

The game objective presented to the player is “*to maintain the number of animals and overall agricultural yield of your choice*.” Thus, the player is asked to make management decisions reflecting their preference of animal population and agricultural yield trajectory. At the end of a game session the player is presented with two scores which allows them to assess their performance relative to their own previous game sessions as well as those of other players. The scores are arbitrarily defined to reflect performance in terms of the animal population (“animal score”) on the one hand, and overall agricultural yield (“yield score”) on the other. Both scores can be interpreted as the mean % of the initial starting score, which is set at 100% (see S1 for further details). They are updated and displayed at each time step, and the final scores are displayed on a score board after the final time step is complete, or once the animal population goes extinct. The scoreboard is a top 10 “leaderboard” of scores over all sessions played by all players to date; if the current player’s score is not included in the top 10, it is displayed at the bottom of the board with the correct rank relative to other players.

#### 3.2.4 Data collection & database

Game play data (e.g. session variables, player inputs, environment state variables) are stored in a MySQL relational database. Database structure is outlined in S1. A full list of parameter values stored, and their description, is given in Table S2. This represents only a subset of all GMSE parameters and may be easily extended in the future by adding additional fields to the relevant database table and ensuring the database interface functions append the extra parameters. For any GMSE parameters that are not stored currently, the default GMSE parameter values are used.

## 4 Example application

### 4.1 “Sandbox” for *in silico* experiments

The combination of the underlying modelling framework, game interface, and the database back-end, provides a platform to collect data on player interaction with the models in a range of simulated environments. This might include *in silico* tests of the effect of specific variability in the environment on simulated animal population extinction, or collecting “big data” on player decision-making given a set of (more or less) variable parameters in terms of animal population, observation, or user models. For example, a researcher using the platform may be interested in testing how human decision-making varies depending on the extent of observed variation in either the ecological (e.g., more or less uncertainty in animal population trajectories) or social (e.g., higher or lower variability in land ownership or sizes of farmer budgets) parts of the modelled system. Data from such experiments may then be combined with debriefing interviews with players to further investigate what may drive such decision-making (e.g., Rakotonarivo, Bell, et al. 2021). Alternatively, by collating large quantities of decision-making data under varying parameter settings, in addition to the outcome of each game session (e.g., animal population extinction and/or trajectories), it may be possible to develop algorithms that can make decisions that are most likely to lead to a desired outcome (e.g. minimising extinction probability while maintaining agricultural yield, or maximising one or the other). While the genetic algorithm for manager decision-making currently implemented in GMSE is effective, it does not currently balance multiple objectives, nor does it necessarily accurately reflect variability in real-life decision-making processes. Parameterising an alternative algorithm based directly on empirical decision-making data has the potential to address these shortcomings.

### 4.2 Example scenario & method

#### 4.2.1 Rationale & methods

Here we illustrate one aspect of this potential by collecting decision-making data from a sample of test players. The main aim was to (1) obtain feedback on the model-game set up, and (2) collect example data to illustrate the potential of the approach, with specific emphasis on how communication of it may be improved in the future. We circulated a link to the game with scenarios configured as detailed below to a sample of 45 contacts working in conservation science and practical conservation and management, covering a range of academic institutions, research institutes, NGOs and government. Contacts were also asked to share the link with any potentially interested contacts. An accompanying covering letter explained this aim, the background to the work, and a request to respond with any feedback. It should be stressed that the data collected here should not be interpreted as comprehensive research on a specific question. It is intended as illustrative of the approach only.

For this proof of concept, we chose to focus on a scenario that systematically varies two parameters, farmer land ownership distribution □_□_ and the number of farmers (□). While inequity in land ownership is commonplace and of interest to conservation strategies (e.g., Rakotonarivo, Bell, et al. 2021), the current manager decision-making algorithm implemented in GMSE cannot explicitly account for the extent of such variation. Thus, collecting empirical data on how decisions and resultant population trajectories may be affected by variable land distribution is important.

Each new game session is initialised with a random draw of one of three possible values of □_□_, representing low, moderate, and high variability in land ownership (resulting landscape patterns illustrated in Figure 3) and one of nine possible values of □, i.e. 4-12 farmers. In addition to this variability, each session also has a small amount of random population mortality (*0.05* ≤ □_□_ ≤ *0.2*), sampled from a uniform distribution. All other parameters are kept constant within and between all game sessions. Although the landscape ownership distribution is clearly shown to the player throughout the game (Figure 2), the player is not told explicitly that ownership will vary before a session starts, or what the extent of this variability will be. This was done to ensure that a player would not selectively abort sessions. Other than this scenario-based parameter variation, game play progresses as described above, with the player able to make management decisions (setting costs for farmer actions) over 20 time steps following the initial five.

**Figure 3.**
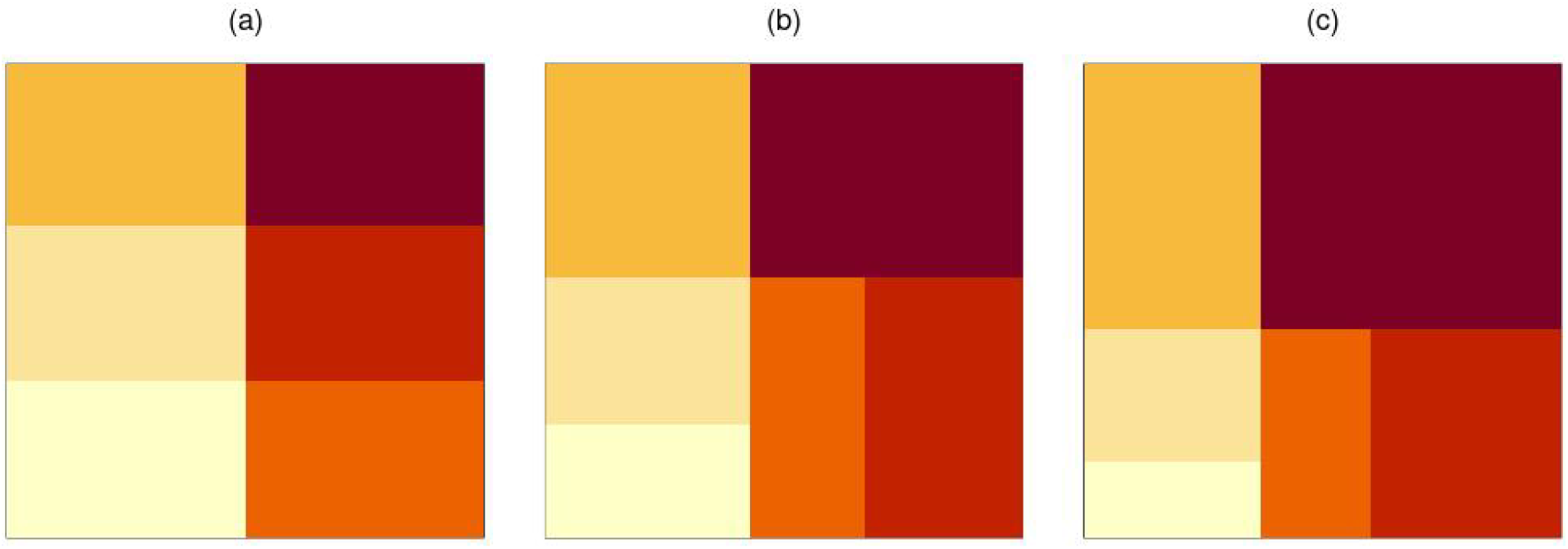
Examples of landscape ownership distributions, (a) low variability, □_□_ = 0 (equal distribution), (b) medium variability, □_□_ = 0.25, and (c) high variability, □_□_ = 0.5, here shown for 6 farmer-landowners.

#### 4.2.2 Ethics

The work described here was approved by the University of Stirling’s General University Ethics Panel (GUEP), project no. 2519. While the game link is publicly accessible, it was not publicised beyond the direct contacts described above. On accessing the link, players are presented with a series of introductory screens explaining the background and purpose of the game, followed by a digital consent form, with a confirmation tick box. No personally identifiable data are collected or stored, other than a player nickname - the latter is only requested so that scores can be shown in context and compared to other players; however this can be left as a default placeholder, and players explicitly told that this is not expected to be their real name. Player nicknames are replaced by random identifiers prior to further data processing.

### 4.3 Illustrative results

Note that the results presented here are intended as illustrative of the model-game approach only, and should be interpreted as such.

Between 21 July 2021 and 19 August 2021, we collated data on 76 play sessions by 28 unique players5; this equated to a total of 1189 decisions (costs set). Sessions lasted 4.5 minutes on average (median 1.6 minutes, range 0.2 - 179.4; the latter maximum duration recorded was an outlier, likely caused by a game session not having been finished and the browser window left open). As per the scenario set up, these sessions were roughly equally distributed between land ownership variability □_□_ (0, 0.25 or 0.5, N = 21 [28%], 32 [42%], and 23 [30%], respectively) and number of farmers □ (4-12).

The animal population reached extinction in 23 out of the 76 sessions (30.3%). Extinction probability appeared to be higher at both higher levels of land ownership variability (□_□_ = *0.25* and □_□_ = *0.5*), particularly so at intermediate (□_□_ = *0.25*) levels (Figure 4a). Differences in extinction probability with variability in farmer (stakeholder) number was less pronounced (Figure 4b).

**Figure 4.**
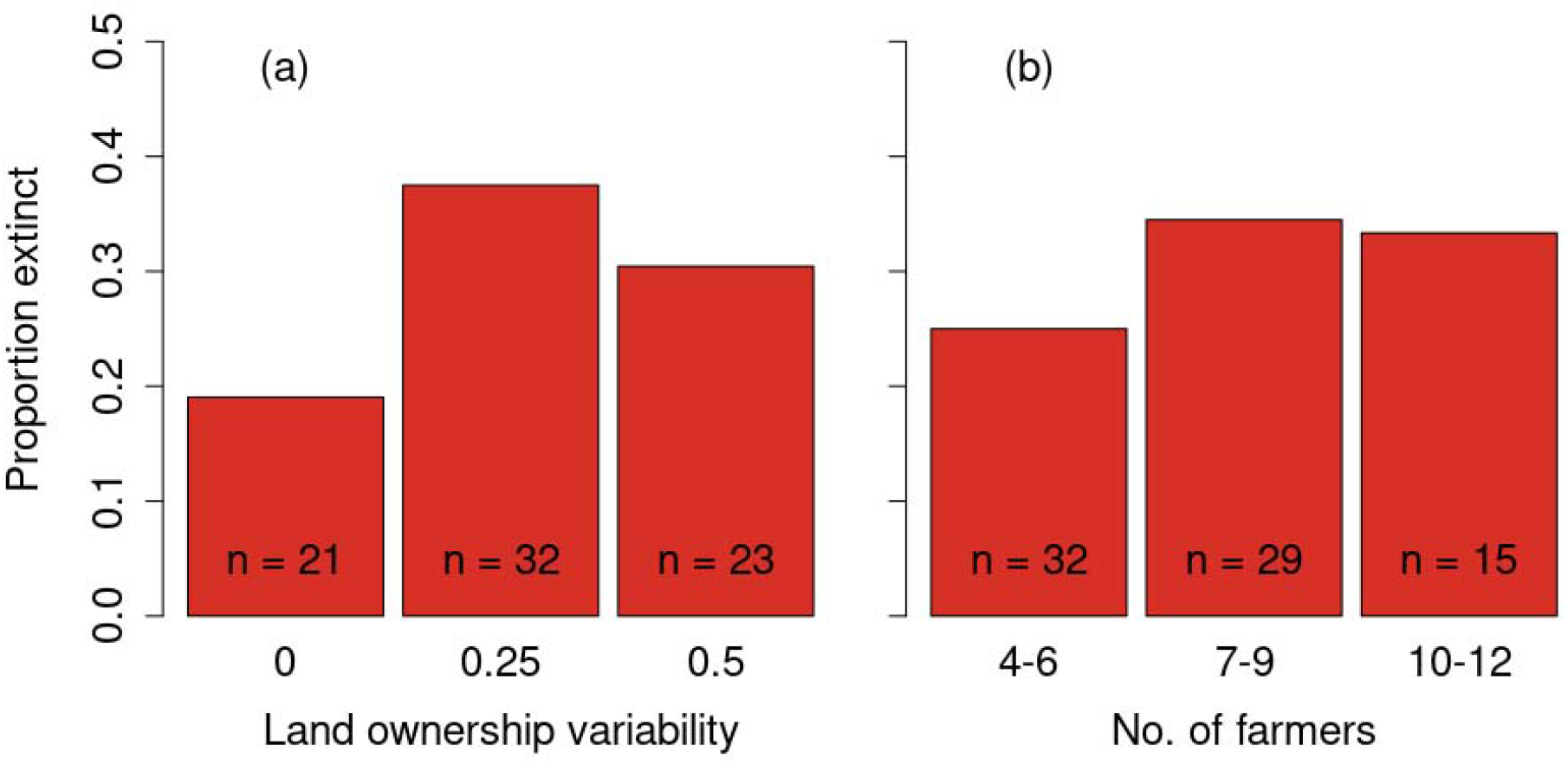
Proportion of game sessions where animal population reached extinction, as a function of (a) land ownership variability and (b) the number of farmers (stakeholders) in the game session.

These extinction probabilities were reflected in the animal population trajectories in each parameter scenario. Figure 5 shows trajectories per level of landownership variability, with cases where the population reached extinction highlighted in red. Both higher levels of variability (□_□_ = *0.25* and □_□_ = *0.5*) show fewer cases with rapid increasing trends.

**Figure 5.**
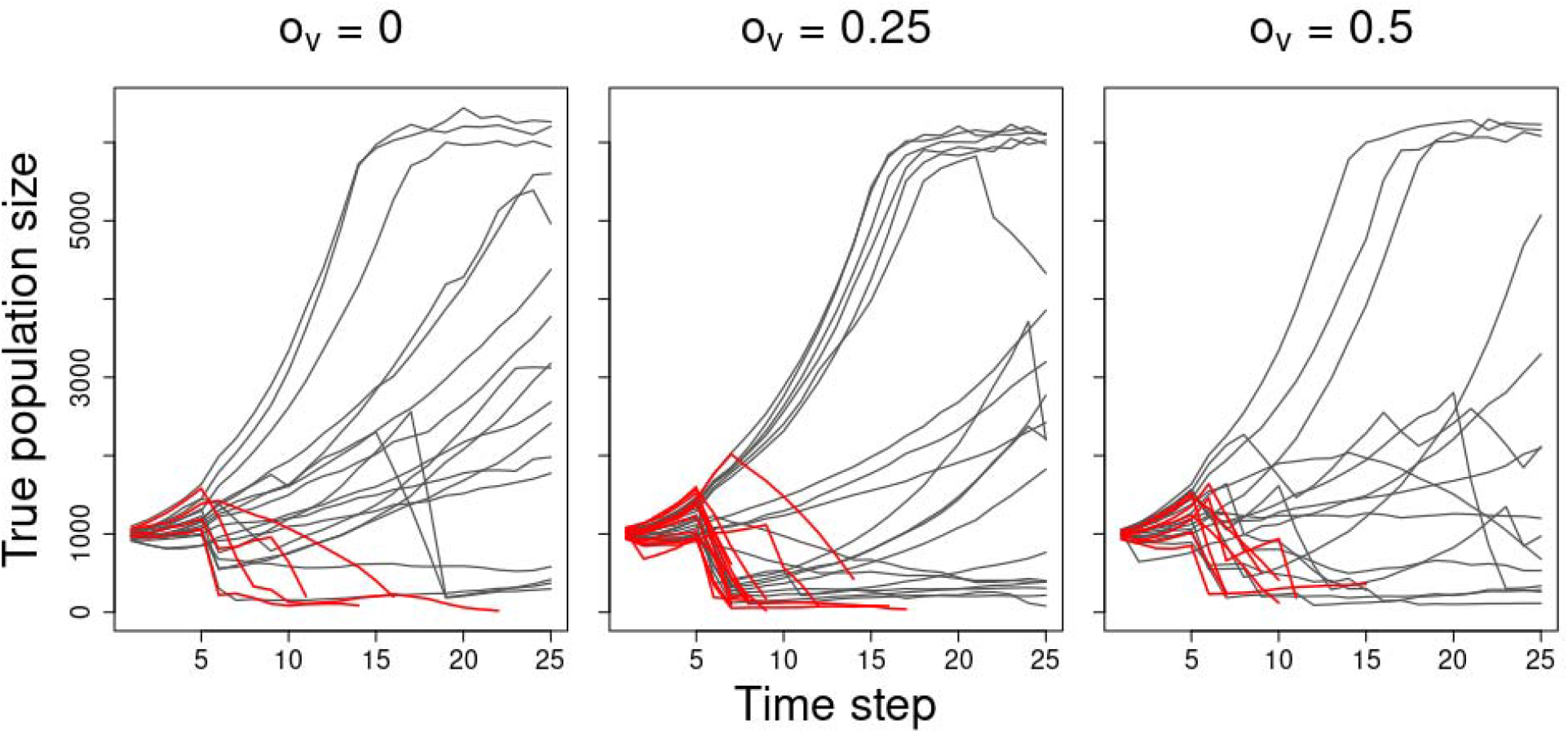
Animal population trajectories per game session, split by levels of land ownership variability. Trajectories highlighted in red are sessions where the population reached extinction.

Management actions taken by the players (over time, □ > *5*) are summarised in Figure 6. It is notable that when land ownership variability was higher (□_□_ = *0.5*), chosen costs for hunting licences appeared to be more stable (i.e., less variable), particularly toward the end of playing sessions (Figures 6c vs. 6a-b). It should be noted that this may in part be an artifact of somewhat lower sample size at higher time steps (because in some sessions the population would have gone extinct part way through a session). On average, hunting licence costs also appeared to be set lower overall at higher land ownership variability. By comparison, costs set for scaring licences appeared to more stable over time (Figures 6d-f).

**Figure 6.**
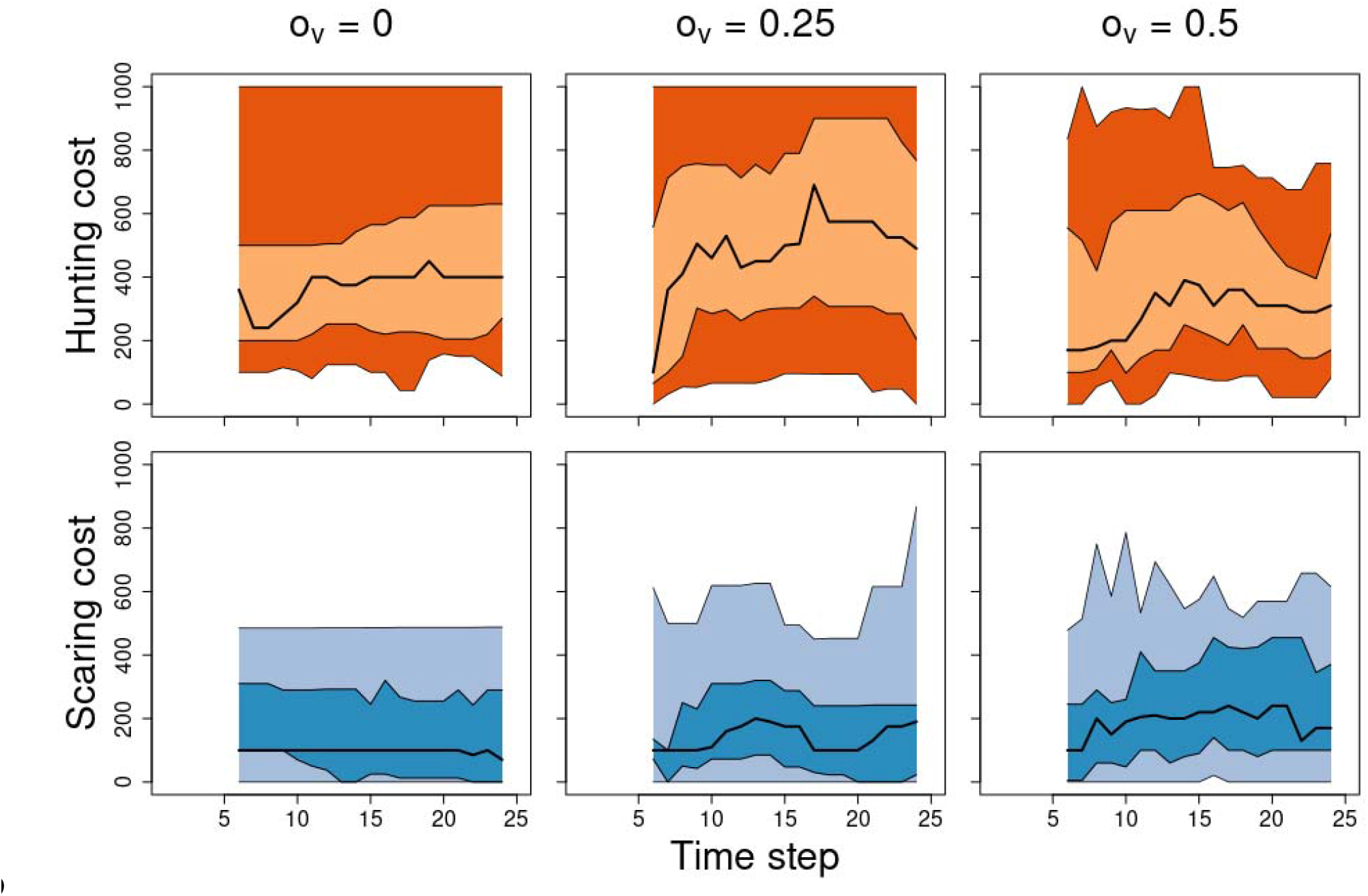
Summary of player management actions (costs set for hunting- and scaring licences) over time, per ownership variability scenario. Solid black line is the mean cost per time step, with lighter and darker polygons representing the 25-75% and 2.5% and 97.5% quantiles of the cost distribution per time step.

## 5 Discussion

We have here outlined a framework for using an interactive game (A&F) as an interface to a social-ecological model for natural resource management. The game interface allows players that are not familiar with the underlying model to interact directly and easily with it, with game play decisions directly reflecting parameter settings in the models. We argue that not only does this provide a convenient communication/education tool with respect to both the specific model and models in general, it also provides a tool to both perform *in silico* experiments on human decision-making in given natural resource management scenarios, as well as collect large amounts of data that may be used to improve the model parameterisation. It is worth noting that we are here specifically referring to model-games as data-collection tools, as opposed to exclusively as communication- or educational tools.

### 5.1 Potential

We illustrated the potential of this approach by presenting data from a small number of trial game play sessions: we showed that subtle variation in farmer land ownership can lead to noticeably different resource population trajectories and manager (player) decision-making patterns. While the data shown here should be taken as illustrative only, it highlights the potential to easily run a range of *in silico* experiments with direct relevance to real-world questions. For example, uncertainty in the estimation of population numbers (observation uncertainty), and its consequences on decision-making is a perennial topic in conservation management (Nuno, Bunnefeld, and Milner-Gulland 2013). While real-world experiments on this would be extremely challenging and costly, GMSE is a suitable modelling framework in which observation uncertainty can be manipulated, and A&F provides a platform to run controlled experiments with real-world stakeholders. This approach could extend to many if not all of the 74 parameters currently controllable by users of GMSE, ranging from variability in demography or behaviour of the natural resource, to farmer behaviour or variability, and wider environmental change or stochasticity. The game interface and player interaction would remain the same, with only the underlying architecture and database back end requiring minor adjustment to accommodate the extra parameter variation.

In addition to use as an experimental tool, this approach also has great potential for use as a way to source large amounts of decision-making data which may then be used to re-parameterise the underlying models, to better reflect real-world decision making. Given a large enough sample of play sessions with a range of parameter combinations and outcomes, it may be possible to train machine learning algorithms on data collected from this approach, to simulate human decision-making under a wide range of conditions (Chabris 2017; Duthie et al. 2021). Such algorithms would potentially reflect a range of subtleties of the decision-making process, including balancing multiple objectives in the presence of competing social, financial, and organisational constraints. Algorithms implemented in existing modelling approaches (without reference to empirical data) including GMSE, are limited in how they can represent such “non-rational” decision-making (Constantino et al. 2021; Dobson et al. 2019).

### 5.2 Some limitations and potential solutions

#### 5.2.1 “The game is unrealistic”

There are a number of limitations to the model-game approach, particularly in terms of directly using “game-sourced” data to (re)parameterise underlying models. One concern raised by several trial players can be summarised as the game or game play lacking “realism,” e.g., lacking aspects or features or real life, or the player’s experience of the conservation problem. This could be particularly problematic if any data collected was subsequently used to adjust model parameterisation: if the game world is not seen as sufficiently realistic, it may be argued that player behaviour is not realistic (i.e. perceived lack of realism leading to lack of external validity; (Jackson 2012; Levitt and List 2007), and therefore any reparameterisation would be biased. While a very important point, it is interesting to note that it relates to the *underlying model* as opposed to the game or the game interface itself. That is, concerns about the lack of “features” or assumptions made are as applicable to any model as they are to the game representation of it, and indeed they are applicable to all models (“*all models are wrong*,” Box 1979). Indeed, this in itself highlights the value of the model-game approach, in that it helps the player to fully understand the model’s structure, assumptions, and consequent limitations: particularly given complex social-ecological models, it can be challenging to effectively communicate the full scope of features and limitations (Grimm et al. 2006, 2020). By casting the model as a game, players are put in the center of the modelling process, and any limitations are likely more apparent, more quickly. Recognition of this, particularly by those lacking technical modelling expertise is vital when such models are put to applied use: all models are abstractions of reality and their utility (“*some models are useful*,” Box 1979) depends on careful application and recognition of this.

#### 5.2.2 “Humans are biased”

An additional limitation of “gamesourcing” data, either in experimental settings or for parameterising models, is the potential for bias in the audience sample. For example, whether intentionally or unintentionally, it may be that players are sampled from a limited subset. All players may have a single professional background such as conservation science, or the nature of the game (framing) may selectively attract a subset of the public (as in known return bias in questionnaires, e.g. Cheung et al. 2017). As a consequence, the in-game decision-making may not be representative of a wider population of potential players, perhaps by the audience in question being more biased towards conservation rather than social objectives. While this is an important potential issue, we argue that it can be avoided by carefully controlling player recruitment, and subsampling of data collected in different sampling regimes, depending on the research question. This may be achieved, for example, by using game play session codes, separating game sessions for a specific experiment from “open” play sessions (Jones et al. *in prep*).

Similar bias may occur if some players play the game with widely different motivations (e.g., Levitt and List 2007), such as playing to “win” versus deliberately attempting to achieve undesirable outcomes. Indeed, it should be stressed that the scores used in the example implementation presented here are to some extent entirely arbitrary, and the choice of scoring system (including algorithms to calculate them) may inherently bias the decision-making data collected. There are a number of ways in which this issue can be addressed. First, when fully implementing this model-game approach, it will be vital to also collect player data through pre- or post-game questionnaires, including on for example professional background and social- and ecological attitudes (as in e.g., Rakotonarivo, Bell, et al. 2021; Rakotonarivo, Jones, et al. 2021), which can be used to control for any potential motivational biases in decision-making data. It should be noted that the current example implementation of A&F allows for anonymous play, and that collection of player personal data would require both further ethical approval as well as additional infrastructure (i.e., unique player names through codes or accounts). Second, it should be stressed that in setting up A&F, we were careful not to steer players to maximise any specific objective (See 3.2.3 above; the goal stated in the introductory screens is “*your aim is to maintain the number of animals and overall agricultural yield of your choice*”). Careful framing of the game objectives (either in open play or in more limited experimental settings) is vital to avoid goal bias (cf. Baynham-Herd et al. 2020).

### 5.3 Conclusions & future direction

Provided that the limitations outlined above are taken into account, and the application is carefully considered, we believe that the approach outlined here has great potential to advance both the understanding and capability of complex social-ecological models for natural resource management. As previous work has already shown, games, and in particular videogames, provide a great tool to increase public engagement with quantitative models, and here we highlight how this could be extended to provide effective, flexible and powerful tools for data collection.

It is worth stressing that the specific parameterisation of the game presented here, as well as the data collected, is intended as illustrative only. The current game could easily be expanded to give the player more control over the game “world” as is required for a given research question, provided it is supported by the underlying model. More broadly, this proof of concept further supports the case for much wider model-game developments (Duthie et al. 2021). Within the broad theme of natural resource management, more sophisticated games might involve “open worlds” in which a plethora of decisions and strategies are available to players, situated in rich environments that may be affected by (and respond to) decisions in a variety of ways - including potentially other decision-makers in multiplayer settings. Many hugely successful modern commercial videogames (e.g. Red Dead Redemption, MineCraft, Sim City) already provide such highly sophisticated environmental simulations, and the potential for sourcing data on human decision-making (and more broadly) from such virtual environments (or similar custom platforms) is huge. Yet, despite recent developments (Crowley, Silk, and Crowley), this potential remains almost untapped in conservation science and natural resource management.

## 6 Acknowledgements

We thank all the trial players for their time and effort in testing A&F. Special thanks to Robbie Whytock for comments on the draft manuscript, and to seven of the trial players for providing specific feedback on which much of the Discussion for this paper was based, and which will form a starting point for future improvements of the model-game approach. JM, LTW and NB were funded by EU Horizon 2020 grant agreement no. 679651 (“ConFooBio,” to NB); ABD was funded by a Leverhulme Trust ECF fellowship grant; MN was funded by the Natural Environment Research Council (IAPETUS Doctoral Training Partnership); IJ was supported by a UKRI Future Leaders Fellowship (MR/T019018/1); AB was funded by IAPETUS NPIF allocation, grant code NE/R012253/1 and supported by NERC.

## Footnotes

1 We here use the term “model” to refer to any predictive quantitative model, although our focus is on predictive simulation models used for decision support. However, the arguments presented here equally apply to statistical models, particularly when used for prediction of trends.

2 Note that there are limitations to this, and that data on decisions made would only be relevant to the context of the game; we discuss limitations in more detail below.

3 A&F currently focuses only on hunting animals, scaring animals or tending crops as available actions to farmers; this may be expanded in the future to other actions available in GMSE.

4 It can be affected *indirectly* by setting the cost for the two actions prohibitively high, so that tending crops becomes more likely to be most beneficial to maximising yield (the farmer’s goal).

5 Strictly speaking, unique player *names*. It is possible for the same player to play under multiple different player names. See Discussion for further details.

