## Supplementary Material 1 for "Models as games: a novel approach for ‘gamesourcing’ parameter data and communicating complex models"

### Supplementary Material 1: Animal & Farm technical details and database structure

#### 1 Animal & Farm technical details

##### 1.1 Structure as relating to GMSE

A&F uses a development version of GMSE implemented in R version 4.1.1 (2021-08-10), the code for which is freely available: https://github.com/ConFooBio/gmse/tree/man_control. In this version, the management model is replaced by player inputs, and the order of operations is altered to accommodate this.

To initialise each game session, four time steps are run using the default GMSE implementation; i.e. in these time steps the management decisions are chosen by the default GA, and the resource, observation and user models are run using the parameters as defined for the given scenario (see 4.2 below). These time steps are followed by a "partial" time step where only the resource and observation models are run, skipping the management and user models. As a result, at the end of these initial time steps (`init_man_control()`), the simulated system has five population and observation time steps and is ready for the next choice of management action at $t=5$, pending the first player input. This is done both to set up all the required GMSE data structures using existing code, as well as to provide the player with a short time series on which to base management decisions going forward.

##### 1.2 Score calculation

The scores are arbitrarily defined to reflect performance in terms of the animal population ("animal score," $A_t$) on the one hand, and overall agricultural yield ("yield score," $S_t$) on the other. Both scores can be interpreted as the mean % of the initial (i.e. at time $t=5$) animal population size and landscape yield, respectively:

$A_t = \frac{1}{N}\sum_{t=6}^{N} n_t \left(\frac{1}{n_6}\right) \cdot 100$ and $S_t = \frac{1}{N}\sum_{t=6}^{N} y_t$

where $t \geq 5$, and $n_t$ is the true size of the animal population at time step $t$, $N$ the total number of animal counts over all time steps in the session, and $y_t$ the mean landscape cell yield over the entire landscape. These calculations are done by the app function `addScores()`.

Both scores are intitialised as $A_t = S_t = 100$ when the game is first started, to ensure score development over a session can be interpreted as a change from that baseline.

#### 1.3 Database backend

In summary, six main tables are used to store data. Tables are linked by the unique session ID present in each table.

- `run` (Table S1a)

  Holding player name, start- and end times for the session and a flag for whether or not the animal population reached extinction or not (single record per session).

- `run_par` (Table S1b)

  Holding game parameters for the GMSE simulation for a session, i.e. all values listed in Table S2. In the example application presented in the main text, the majority of the parameters in this table will be constants, with only `ownership_var` and `remove_pr` varying per session. Note that any GMSE parameters not stored in this table or listed in Table S2 are kept as the default in GMSE, which can be found in the `GMSE` package reference manual (https://cran.r-project.org/web/packages/GMSE/GMSE.pdf) listed under the function `gmse()`.

- `scores` (Table S1c)

  Holding the number of time steps achieved per session and the animal and yield score.

- `gdata` (Table S1d)

A record per time step for each session, recording the true and observed population state, the number of actions of each type taken, and the costs set by the manager (player), as well as the total yield in the environment.

- `yield` (Table S1e)

  The yield achieved by each farmer in each time step, per session.

Records in tables `run`, `run_par` and `scores` are only updated at the start and end of each session, whereas `gdata` and `yield` are the “live” tables that are appended to at each time step during a game session. End times are recorded for each session where the player either reaches $t = 25$, manually resets the game during a session, or as the animal population reaches extinction; i.e. when this field remains blank (`NULL`), it means that a session was not terminated “normally,” i.e. by the browser being closed manually or timing out due to inactivity.

*Table S1(a). Description of database table* `run`*. Unique line per game session ID. 'Type' is the MySQL variable type and NULL indicates whether the value is allowed to be empty.*

| Field | Type | Null | Description |
|---|---|---|---|
| id | INT | NO | Unique game session ID. Acts as primary key to other tables. |
| session | VARCHAR(32) | NO | Internal session string as genrated by the Shiny server. |
| player | VARCHAR(64) | YES | Player name as input. |
| startTime | VARCHAR(23) | NO | Date-time stamp for when the session was started. |
| endTime | VARCHAR(23) | YES | Date-time stamp for when the session was manually ended, the animal population reached extinction, or the maximum number of time steps was reached. If this value is blank, it means the session was not ended by any of these means (e.g. browser closed or crashed). |
| extinct | TINYINT(1) | NO | A flag indicating whether the animal population reached extinction during the session. |

*Table S1(b). Description of database table* `run_par`*. Unique line per game session ID, storing GMSE parameter values for session. 'Type' is the MySQL variable type and NULL indicates whether the value is allowed to be empty.*

| Field | Type | Null | Description |
|---|---|---|---|
| id | INT | NO | Unique game session ID. Acts as primary key to other tables. |
| K | INT | YES | Number of initial time steps (before player input starts) |
| land_ownership | TINYINT(1) | YES | See Table S2 |
| stakeholders | INT | YES | See Table S2 |
| manager_budget | INT | YES | See Table S2 |
| manage_target | INT | YES | See Table S2 |
| observe_type | INT | YES | See Table S2 |
| res_move_obs | INT | YES | See Table S2 |
| res_death_k | INT | YES | See Table S2 |
| lambda | FLOAT | YES | See Table S2 |
| res_death_type | INT | YES | See Table S2 |
| remove_pr | FLOAT | YES | See Table S2 |
| user_budget | INT | YES | See Table S2 |
| culling | TINYINT(1) | YES | See Table S2 |
| scaring | TINYINT(1) | YES | See Table S2 |

| Field | Type | Null | Description |
|---|---|---|---|
| tend_crops | TINYINT(1) | YES | See Table S2 |
| land_dim_1 | INT | YES | See Table S2 |
| land_dim_2 | INT | YES | See Table S2 |
| resource_ini | INT | YES | See Table S2 |
| tend_crop_yld | FLOAT | YES | See Table S2 |
| max_years | INT | YES | Maximum number of time steps for player |
| public_land | FLOAT | YES | See Table S2 |
| ownership_var | FLOAT | YES | See Table S2 |
| usr_budget_rng | INT | YES | Farmer budget range. Not used in example implementation. |
| score_display | CHAR(5) | YES | Identifier indicating which score display was used; internal and testing use only. |

*Table S1(c). Description of database table* `scores`*. Unique line per game session ID, storing session score data. 'Type' is the MySQL variable type and NULL indicates whether the value is allowed to be empty.*

| Field | Type | Null | Description |
|---|---|---|---|
| id | int | NO | Unique game session ID. Acts as primary key to other tables. |
| steps | int | YES | Number of game time steps achieved |
| mean_res | int | YES | Animal score for session |
| mean_yield | int | YES | Yield score for session |
| total | int | YES | Total (combined) score for session (not used) |
| score_version | int | YES | Indicator for score version calculation. Internal and testing use only. |

*Table S1(d). Description of database table* `gdata`*. Unique line per game session and time step; stores observations (of animal population), farmer actions, and player actions per time step. 'Type' is the MySQL variable type and NULL indicates whether the value is allowed to be empty.*

| Field | Type | Null | Description |
|---|---|---|---|
| id | INT | NO | Unique game session ID. Acts as primary key to other tables. |
| t | INT | YES | Time step |
| res | INT | YES | True number of animals |
| obs | FLOAT | YES | Observed number of animals |
| culls | INT | YES | Number of hunting actions |
| scares | INT | YES | Number of scaring actions |
| tend_crops | INT | YES | Number of crop tending actions |
| cull_cost | INT | YES | Hunting cost set for farmers in next time step (blank for final recorded time step for given session, as this is completed in response to user actions in the current time step. |

| Field | Type | Null | Description |
|---|---|---|---|
| scare_cost | INT | YES | Scaring cost set for farmers in next time step (blank for final recorded time step for given session, as this is completed in response to user actions in the current time step. |
| yield | INT | YES | Total landscape yield. Legacy data and not currently used. |

73

*Table S1(e). Description of database table* `yield`*. Unique line per farmer within time step, within session. 'Type' is the MySQL variable type and NULL indicates whether the value is allowed to be empty.*

| Field | Type | Null | Description |
|---|---|---|---|
| id | int | NO | Unique game session ID. Acts as key to other tables. |
| t | int | NO | Time step |
| user | int | NO | Farmer identifier |
| pyield | float | NO | Mean landscape cell yield for given farmer in given time step. |
